# Insights into Human Norovirus Cultivation in Human Intestinal Enteroids

**DOI:** 10.1101/2024.05.24.595764

**Authors:** Khalil Ettayebi, Gurpreet Kaur, Ketki Patil, Janam Dave, B. Vijayalakshmi Ayyar, Victoria R Tenge, Frederick H. Neill, Xi-Lei Zeng, Allison L. Speer, Sara C. Di Rienzi, Robert A. Britton, Sarah E. Blutt, Sue E. Crawford, Sasirekha Ramani, Robert L. Atmar, Mary K. Estes

## Abstract

Human noroviruses (HuNoVs) are a significant cause of epidemic and sporadic acute gastroenteritis worldwide. The lack of a reproducible culture system hindered the study of HuNoV replication and pathogenesis for almost a half-century. This barrier was overcome with our successful cultivation of multiple HuNoV strains in human intestinal enteroids (HIEs), which has significantly advanced HuNoV research. We optimized culture media conditions and generated genetically-modified HIE cultures to enhance HuNoV replication in HIEs. Building upon these achievements, we now present new insights to this culture system, which involve testing different media, unique HIE lines, and additional virus strains. HuNoV infectivity was evaluated and compared in new HIE models, including HIEs generated from different intestinal segments of individual adult organ donors, HIEs from human intestinal organoids produced from directed differentiation of human embryonic stem cells into intestinal organoids that were transplanted and matured in mice before making enteroids (H9tHIEs), genetically-engineered (J4*FUT2* knock-in [*KI*], J2*STAT1* knock-out [*KO*]) HIEs, as well as HIEs derived from a patient with common variable immunodeficiency (CVID) and from infants. Our findings reveal that small intestinal HIEs, but not colonoids, from adults, H9tHIEs, HIEs from a CVID patient, and HIEs from infants support HuNoV replication with segment and strain-specific differences in viral infection. J4*FUT2-KI* HIEs exhibit the highest susceptibility to HuNoV infection, allowing the cultivation of a broader range of GI and GII HuNoV strains than previously reported. Overall, these results contribute to a deeper understanding of HuNoVs and highlight the transformative potential of HIE cultures in HuNoV research.

**Importance:** HuNoVs cause global diarrheal illness and chronic infections in immunocompromised patients. This manuscript reports approaches for cultivating HuNoVs in secretor positive human intestinal enteroids (HIEs). HuNoV infectivity was compared in new HIE models, including ones from i) different intestinal segments of single donors, ii) human embryonic stem cell-derived organoids transplanted into mice, iii) genetically-modified lines, and iv) a patient with chronic variable immunodeficiency disease. HIEs from small intestine, but not colon, support HuNoV replication with donor, segment and strain-specific variations. Unexpectedly, HIEs from one donor are resistant to GII.3 infection. The genetically-modified J4*FUT2-KI* HIEs enable cultivation of a broad range of GI and GII genotypes. New insights into strain-specific differences in HuNoV replication in HIEs support this platform for advancing understanding of HuNoV biology and developing potential therapeutics.

## Introduction

Human noroviruses (HuNoVs) are the most common cause of both epidemic and sporadic acute gastroenteritis worldwide. They also cause chronic infections in immunocompromised cancer, transplant patients, and patients with primary immune deficiencies and are a severe health burden for this population (1–3). Annually, HuNoV infections are responsible for 685 million cases of diarrhea and over 212,000 deaths globally (4). In the United States alone, these infections lead to approximately 109,000 hospitalizations, generating a substantial economic impact of more than $10.6 billion in healthcare expenses and societal costs (4–6). Even though HuNoV was discovered in 1968, there are still no FDA-approved vaccines and efficacious therapeutics available, highlighting the challenges associated with combating these infections and the need for further research and development in this area (7, 8).

One of the main obstacles in studying and understanding HuNoVs was the absence of a culture system to effectively cultivate these viruses. However, in 2016, a significant breakthrough was achieved with the establishment of the human intestinal stem cell-derived enteroid (HIE) system for HuNoV cultivation (9). This development overcame a five-decade barrier in laboratory models for HuNoVs, opening new avenues for investigating HuNoV biology. Since the establishment of the HIE system, significant advancements have been made in unraveling various aspects of HuNoV biology, including strain-specific differences in replication, methods of virus inactivation, innate immune responses, antiviral susceptibility, and characterization of neutralizing antibodies (10–29). Here, we report further advancements in the HIE culture system by testing HuNoV infection in multiple unique HIE lines and expanding the profile of cultivatable HuNoV strains.

## Results

We previously reported that differentiation of HIEs and infection carried out in the commercial IntestiCult™ Organoid Growth Medium (OGMd referred to as INTd in (23)) from STEMCELL Technologies (formerly known as IntestiCult media) supports robust replication of HuNoV strains that replicated poorly in HIEs grown in our original in-house culture medium (23). A new IntestiCult™ Organoid Differentiation Medium (ODM) media (also from STEMCELL Technologies) became available in 2020, and we conducted a comparative analysis between OGM media and the new ODM media to assess their impact on HuNoV replication. For these studies, we used the previously reported genetically-modified J4*FUT2-KI* HIE line in which the fucosyltransferase 2 (*FUT2*) gene was knocked-in to the secretor negative J4 HIE (30). The J4*FUT2-KI* HIE line showed better HuNoV binding to the HIE cell surface and replication of several HuNoVs (30). J4*FUT2-KI* HIE monolayers, seeded in the proliferation OGM medium (OGMp) and then differentiated in either OGMd or ODM media, were infected with a GII.4 Sydney[P31] or a GII.3[P21] HuNoV strain. Both media supported viral replication. However, replication of both strains was enhanced based on significant greater geometric mean log_10_ genome equivalent (GE) increases in virus yields from 1 hour post infection (hpi) to 24 hpi in HIEs differentiated using OGMd media when compared to ODM media (Figure 1). Consequently, all subsequent experiments described in this study were executed using OGM media. Infections were conducted using the OGMd medium supplemented with the bile acid sodium glycochenodeoxycholate (GCDCA), necessary for GII.3 infection and enhancing GII.4 infection (9).

**Figure 1:**
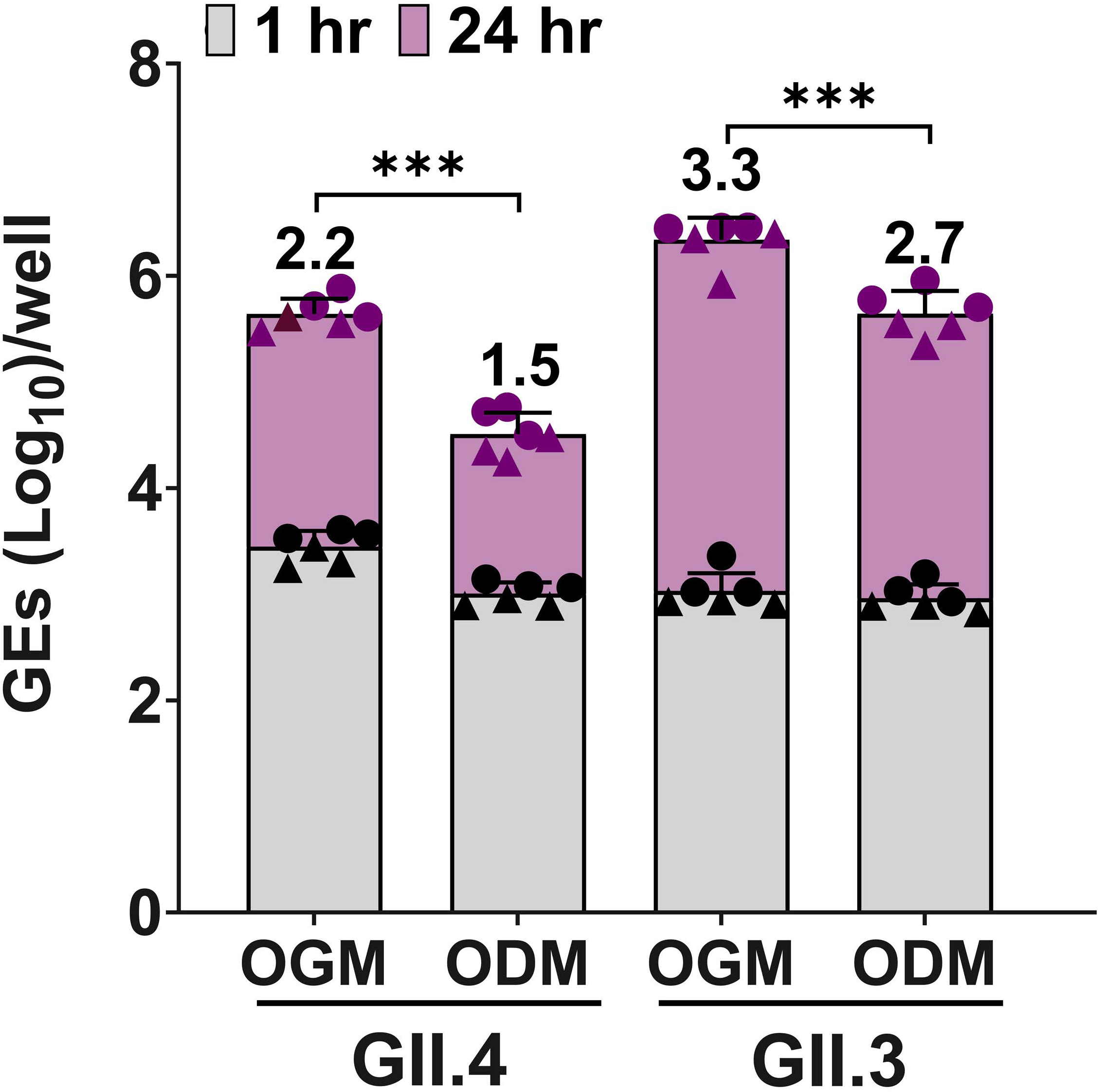
HuNoV replication is enhanced in HIEs plated in OGM compared to ODM media. 3D J4*FUT2-KI* HIEs were propagated in WRNE media. Monolayers were seeded in OGM proliferation medium with ROCK inhibitor and differentiated in OGMd or ODM differentiation media. Differentiated monolayers were inoculated with GII.4 Sydney[P31] (9 × 10^5^ GEs/well) or GII.3[P21] (4.3 × 10^5^ GEs/well) diluted in CMGF(-) with 500 μM GCDCA. After 1 hpi, monolayers were washed twice and cultured in the OGMd or ODM media (+ 500 μM GCDCA). Values above the bars represent net log_10_ difference in viral growth (Δ24hr-1hr). Gray shading indicates GE at 1 hpi and purple shading shows GE at 24 hpi. Mean data compiled from two independent experiments with three wells per experiment are shown; error bars show standard deviation (SD). Experiments are denoted with different symbol shapes (circle or triangle). Significance was determined at 24 hpi using Student’s *t*-test. *** p value < 0.001.

We next conducted a comparative investigation of viral replication using multiple, unique secretor positive HIE lines derived from various sources. We aimed to capture segment-specific and strain-specific differences in the susceptibility of HIE lines that may influence viral replication. We first evaluated HIEs generated from intestinal tissues obtained from four independent organ donors (designated 2002, 2003, 2004, and 2005). From each donor, four HIE lines were established, representing the three segments of the small intestine [duodenum (D), jejunum (J), and ileum (I)] as well as the colon (C). The HIE lines were confirmed to be secretor positive through genotyping assays for fucosyltransferase 2 (*FUT2*), which indicated their ability to express the histoblood group antigens (HBGAs) necessary for HuNoV infection. The susceptibility of HIE lines to HuNoV strains GII.4 Sydney[P31] and GII.3[P21] was assessed, focusing on segment-specific differences (Table 1).

**Table 1:**
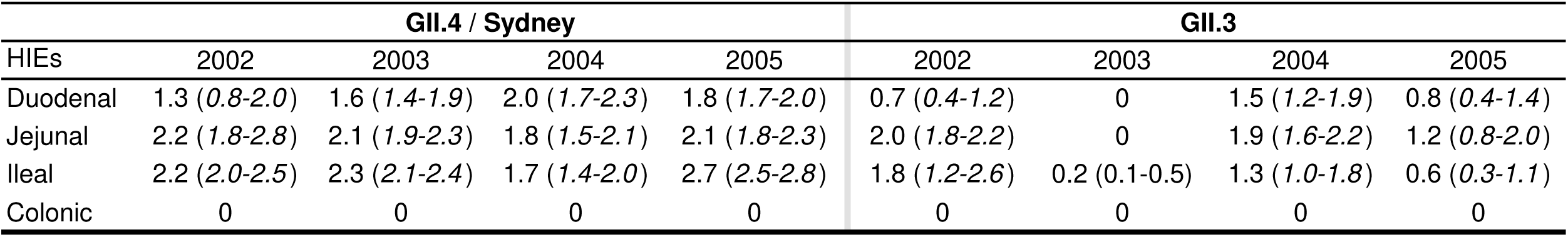
HuNoV yields after replication in HIEs from four donors. Differentiated monolayers were inoculated with GII.4 Sydney[P31] (9 × 10^5^ GEs/well) or GII.3[P21] (4.3 × 10^5^ GEs/well) diluted in CMGF(-) with 500 μM GCDCA. After 1 hpi, monolayers were washed twice and cultured in the OGMd medium (+ 500 μM GCDCA). Values represent compiled data of net log_10_ geometric mean increase in viral growth (Δ24hpi-1hpi) from 2 experiments and 3 replicate wells in each experiment; values in parentheses correspond to 95% confidence interval (95% CI).

GII.4 replication was observed across all small intestinal HIE lines. For GII.4, the yields from the duodenal HIE lines were similar between the lines except 2004 was significantly greater than 2002 (p<0.01). For GII.4, the yields in the jejunal HIE lines were similar across all four lines. The yields for GII.4 in the ileal HIE lines were significantly different from each other (p<0.01) except between the 2002 and 2003 lines.

GII.3 replication occurred in HIE lines generated from small intestinal segments of three out of the four donors. Unexpectedly, GII.3 replication was not detected in any of the HIEs generated from donor 2003. Comparing replication in the other three lines, GII.3 yield in the duodenal 2004 HIE was significantly greater than in 2002 and 2005 (p<0.05) HIEs. In the jejunal lines, GII.3 yield was significantly less in 2005 compared to 2002 (p<0.01) and 2004 (p<0.05). In the ileal lines, GII.3 yield was significantly less in the 2005 line than in the 2002 (p<0.001) and 2004 (p<0.05) lines.

None of the colonic HIE lines supported replication of either GII.3 or GII.4 HuNoV strains. These observations suggest that while small intestinal HIE lines are generally permissive to HuNoV infection, there may be variations in viral replication efficiency within different segments of the small intestine from individual donors but data from more HIE lines are needed for definitive conclusions. The observed striking difference in infectivity between GII.4 and GII.3 HuNoV for HIEs from donor 2003 also indicates strain-specific differences in susceptibility to infection within the same donor that are independent of FUT2 expression. The 2003 line is the first secretor positive HIE line that we have tested that does not support GII.3 replication. Further, these findings confirm our previous report of lack of replication in colonic lines derived from different independent individuals (23), indicating segment-specific differences in the susceptibility of HIE lines to HuNoV infection.

In addition, we assessed HuNoV infection in other unique HIEs. First, we evaluated viral replication in HIEs generated from directed differentiation of H9 human embryonic stem cells (H9hESC) into intestinal organoids (HIOs) that were transplanted and matured in immunocompromised mice for 8 weeks before making enteroids (referred to in this paper as H9tHIEs). The initial HIOs contained both epithelial and mesenchymal cells (31). Previous RNASeq analyses of such HIOs showed these cultures exhibit a more fetal-like transcriptional expression pattern; they are therefore considered immature (32). However, *in vivo* transplantation of such HIOs into mice (tHIOs) induces maturation of the intestinal epithelium resulting in tHIOs with a more adult-like transcriptional profile and enhanced levels of tight junction proteins (32–35). Additionally, the morphology and cellular maturation patterns in tHIOs are similar to fetal intestinal epithelial development (36). For example, an 8-week-old tHIO demonstrates proliferation confined to the crypts similar to a gestational week 18 human fetal intestine (36). To investigate whether matured tHIOs could be susceptible to HuNoV infection, we transplanted immature HIOs under the kidney capsule of immunocompromised mice. After eight weeks, the matured tHIOs were harvested and processed to proliferate into three-dimensional H9tHIEs, following the same protocol used for generating HIEs from human intestinal tissues and biopsies. The resulting H9tHIEs could be passaged indefinitely, and they also are epithelial only cultures without an outer mesenchyme, a morphology similar to tissue stem cell-derived HIEs (37).

We next investigated whether the five-day differentiated H9tHIEs would support replication of GII.4 Sydney[P31] and GII.3[P21] HuNoVs. Efficient replication of both GII genotypes was achieved (Figure 2A and 2B). A direct comparison of virus replication in the H9tHIEs and our commonly used lab prototype J2 HIE, used extensively in many studies (9), and J4*FUT2-KI* HIE lines showed that H9tHIEs supported GII.4 replication with no significant difference in viral yield when compared to the J2 and a significant reduction in viral yield when compared to the J4*FUT2-KI* line. For GII.3, there was a higher viral yield in the H9tHIE line compared to replication in the J2 HIE line, but the best replication was in the J4*FUT2-KI* HIE line. Both HuNoV strains had significant replication increases in J4*FUT-KI* compared to the J2 line.

**Figure 2:**
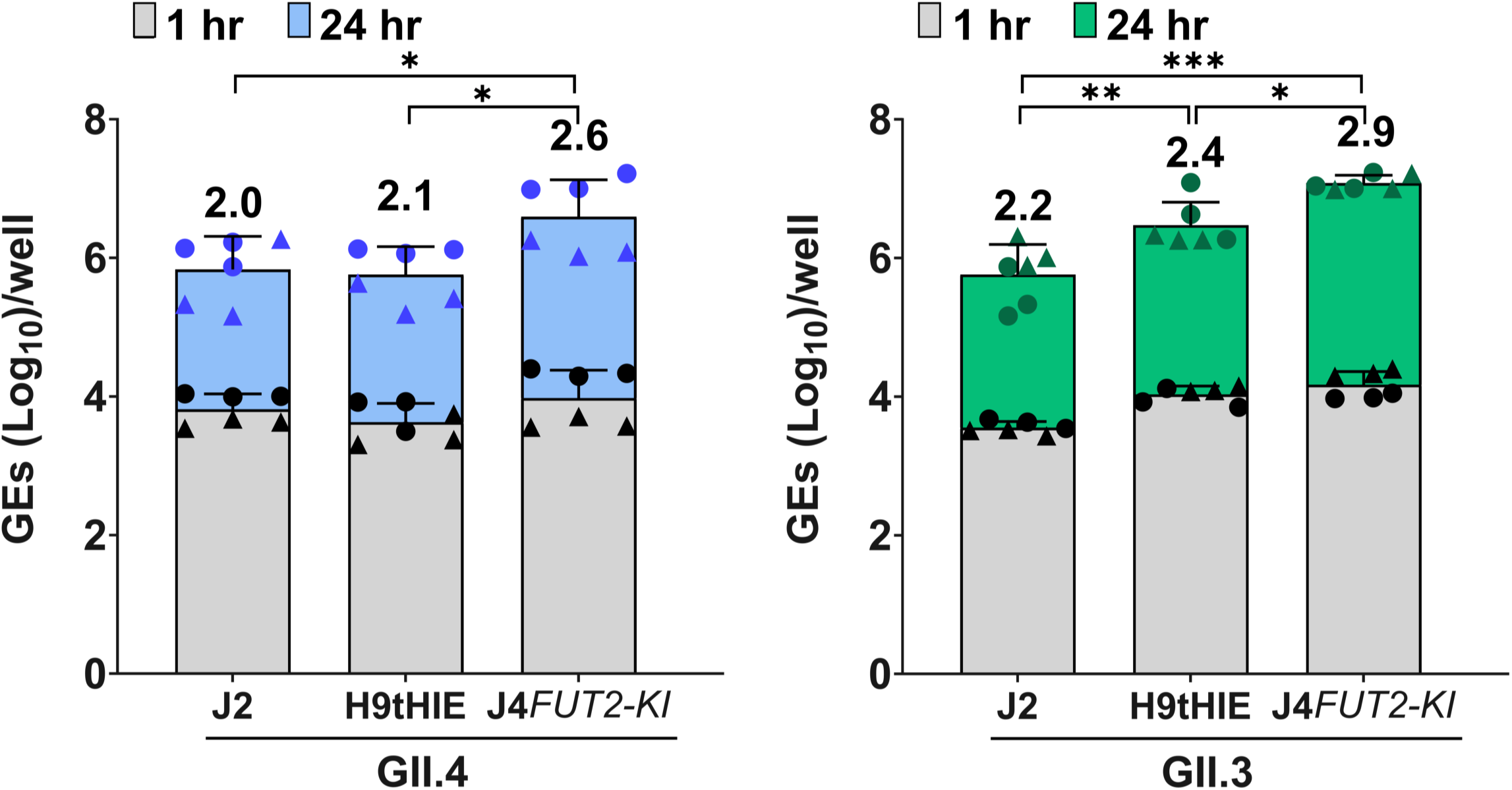
HuNoV replication in H9tHIEs. Differentiated H9tHIE monolayers in OGM differentiation medium were infected with **(A)** GII.4 Sydney[P31] (9 × 10^5^ GEs/well) or **(B)** GII.3[P21] (4.3 × 10^5^ GEs/well) in presence of 500 μM GCDCA. After 1 hpi, monolayers were washed twice and cultured in OGM differentiation medium (+ 500 μM GCDCA). Values on bars represent the net log_10_ difference in viral growth (Δ24hr-1hr). Gray shading indicates GE at 1 hpi and blue (A) or green (B) shows GE at 24 hpi. Mean data compiled from two independent experiments with three wells per experiment are shown; error bars show SD. Experiments are denoted by different symbol shapes (circle or triangle). Significance was determined at 24 hpi. *** p value < 0.001; ** p < 0.01; * p < 0.05.

We next evaluated viral replication in an HIE line (D_201_HIE) generated from a duodenal biopsy obtained from a patient with common variable immunodeficiency disease (CVID) who had been suffering from chronic GII.6 HuNoV infection for years. We assessed the replication of GII.4, GII.3 and two GII.6 isolates in the differentiated D_201_HIE line. The GII.6 virus (BCM18-1) derived from an adult CVID patient failed to replicate in this D_201_HIE line (Figure 3), while the D_201_HIEs supported the replication of GII.4 Sydney[P31] and GII.3[P21] with geometric mean log_10_(GE) increases of 2.7 and 2.3, respectively. The lack of replication of the patient’s own virus was not likely due to the inability of this line to support GII.6 HuNoV replication because a different clinical isolate of GII.6 virus (TCH13-106) from a child hospitalized with acute gastroenteritis showed a minimal replication of 0.5 geometric mean log_10_ (GE) increase (Figure 3).

**Figure 3:**
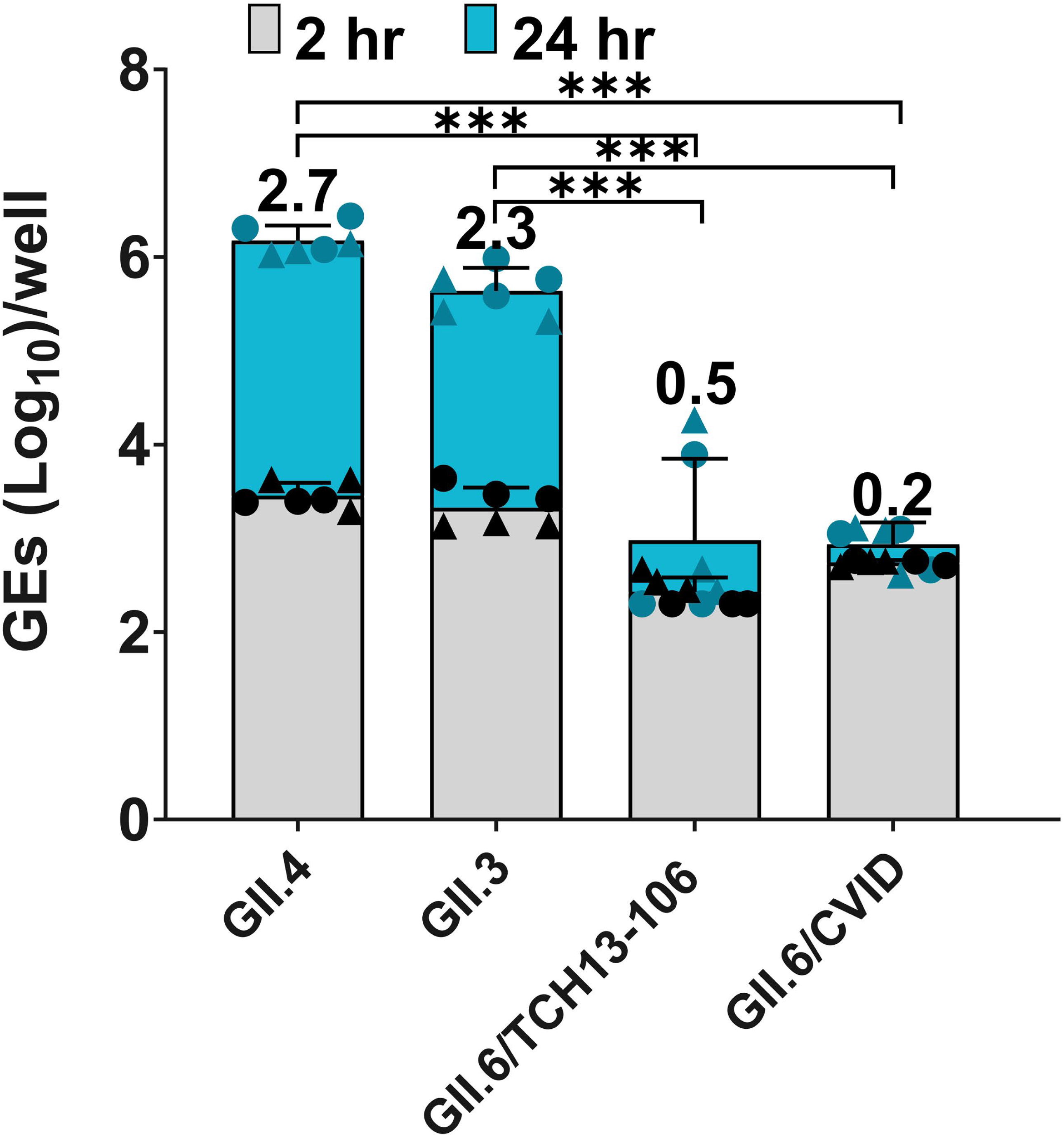
HuNoV replication in D201. D201 HIE monolayers were differentiated in OGM differentiation medium and infected with GII.4/Sydney (9 × 10^5^ GEs/well), GII.3 (4.3 × 10^5^ GEs/well), GII.6/TCH13-106 (3.3 x 10^5^ GEs/well) or GII.6/CVID (BCM18-1) (9.7 x 10^4^ GEs/well) diluted in CMGF(-) supplemented with 500 μM GCDCA. After 2 hpi, monolayers were washed twice and cultured in OGM differentiation medium (+ 500 μM GCDCA). Values on bars represent net log_10_ difference in viral growth (Δ24hr-1hr). Gray shading indicates GE at 2 hpi and blue shows GE at 24 hpi. Mean data compiled from two independent experiments with three wells per experiment are shown; error bars show SD. Experiments are denoted by different symbol shapes (circle or triangle). Significance was determined at 24 hpi. *** p value < 0.001.

We also investigated whether the GII.6 virus inoculum from the CVID patient might not be infectious by testing replication of this virus in three other jejunal HIE lines, including the J2 HIE line and two genetically-modified J2*STAT1-KO* and J4*FUT2-KI* HIE lines made previously (30). The GII.6/CVID patient-derived virus showed minimal or no replication in J2 and J2*STAT1-KO* HIEs, respectively, but it did replicate with a geometric mean log_10_(GE) increase of 1.2 in the J4*FUT2-KI* HIE (Figure 4A). A longer replication time (72 hpi) was evaluated because of the low titer of the CVID virus inoculum. The other GII.6 isolate (TCH13-106), which replicated minimally in the D_201_HIE line (Figure 3) was tested as a positive control and replicated in each jejunal HIE line (Figure 4B).

**Figure 4:**
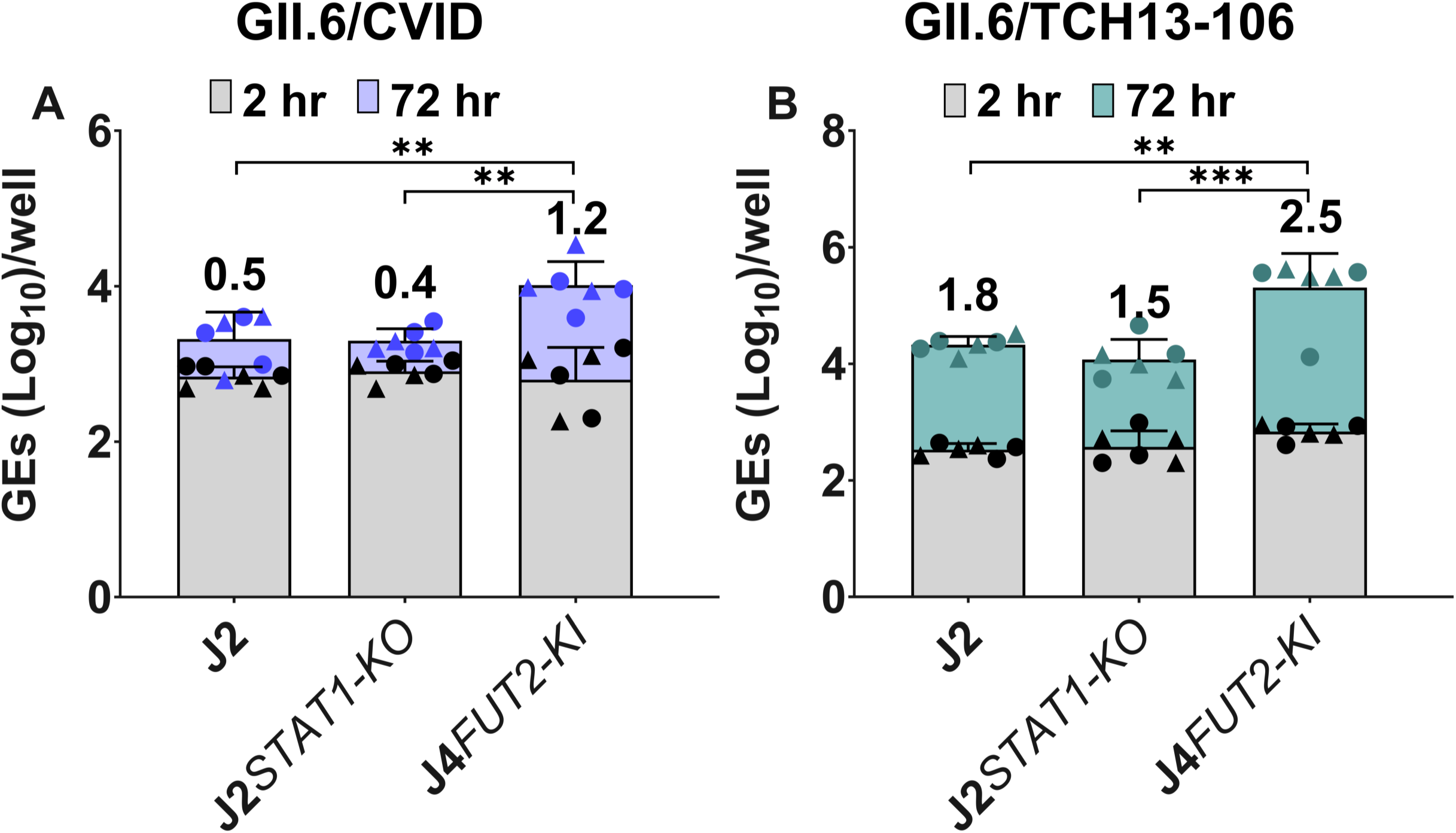
HuNoV replication in genetically modified HIEs. HIE monolayers were differentiated in OGM differentiation medium and inoculated with **(A)** GII.6/CVID (9.7 x 10^4^ GEs/well) or **(B)** GII.6/TCH13-106 (3.3 x 10^5^ GEs/well) in presence of 500 μM GCDCA. After 1-2 hpi, monolayers were washed twice and cultured in OGM differentiation medium (+ 500 μM GCDCA). Values on bars represent net log_10_ difference in viral growth (Δ24hr-1hr). Gray bars indicate GE at 2 hpi and colored bars show GE at 72 hpi. Mean data compiled from two independent experiments with three wells per experiment are shown; error bars show SD. Experiments are denoted by different symbol shapes (circle or triangle). Significance was determined at 24 hpi. *** p value < 0.001; ** p < 0.01.

Next, we evaluated GII.4 and GII.3 viral replication in jejunal HIE lines (J1005 and J1006) established from specimens collected during surgery from two preterm infants, 37 and 43 weeks corrected gestational age at the time of surgical sample collection, respectively (38). Virus replication in these lines was compared to that in the prototype J2 line and the J4*FUT2-KI* HIE line. GII.4 replication in the prototype J2 was similar in the J1005 and J1006 lines. GII.4 replication was significantly higher in J4*FUT2-KI* HIEs compared to the J005 and J006 HIE lines. GII.3 replication was significantly different in J2 or J4*FUT2-KI* between all lines tested with significantly less GII.3 replication in the infant lines compared to the adult lines. Nevertheless, both GII.4 and GII.3 strains replicated in all four HIE lines tested (Figure 5).

**Figure 5:**
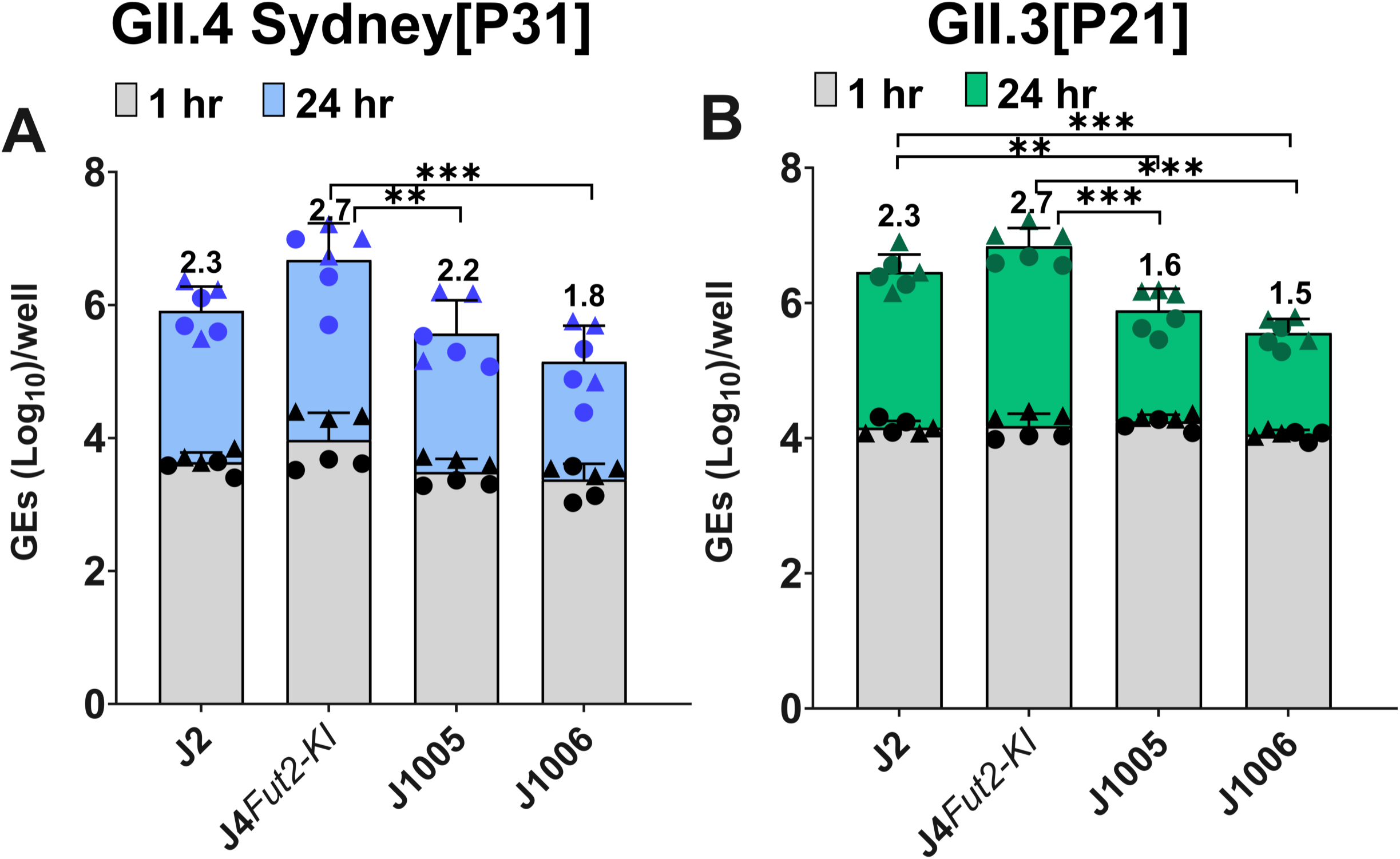
HuNoV replication in HIEs from adults and infants. HIE monolayers were differentiated in OGM differentiation medium and infected with **(A)** GII.4/Sydney (9 × 10^5^ GEs/well) or **(B)** GII.3 (4.3 × 10^5^ GEs/well) diluted in CMGF(-) supplemented with 500 μM GCDCA. After 1 hpi, monolayers were washed twice and cultured in OGM differentiation medium (+ 500 μM GCDCA). Values on bars represent net log_10_ difference in viral growth (Δ24hr-1hr). Gray bars indicate GE at 1 hpi and colored bars show GE at 24 hpi. Mean data compiled from two independent experiments with three wells per experiment are shown; error bars show SD. Experiments are denoted by different symbol shapes (circle or triangle). Significance was determined at 24 hpi with significant differences between groups shown. *** p value < 0.001; ** p < 0.01.

Finally, we evaluated whether these lines exhibit quantitatively different susceptibilities to infection with the GII.4 and GII.3 viruses by determining the number of genomic equivalents per 50% tissue culture infectious dose (TCID_50_) for each virus for each line. The lowest GE/TCID_50_ was seen for the GII.3 virus in the J4*FUT2-KI* HIE [3.5 (*1.01*)], [log_10_(GE)/TCID50, standard deviation (SD)], followed by the infant HIEs [3.6 (*1.09*) and 3.6 (*1.02*) for J1005 and J1006, respectively]. Surprisingly, the GE/TCID_50_ of GII.4 Sydney[P31] HuNoV was significantly higher in the J4*FUT2-KI* line [4.3 (*1.06*)] compared to J2 [3.9 (*1.07*)] and infant HIEs [3.7 (*1.05*) and 3.7 (*1.06*) for J1005 and J1006, respectively] (Table 2). To explore the specificity of this observed high GE/TCID_50_ for GII.4, we tested another GII.4 Sydney strain with a different polymerase type, GII.4 Sydney[P16] (38), in this assay. The GE/TCID_50_ of this strain reflected the pattern observed for GII.4 Sydney[P31], with elevated GE/TCID_50_ values particularly seen in the J4*FUT2-KI* line [5.1 (*1.01*)] (Table 2).

**Table 2:**
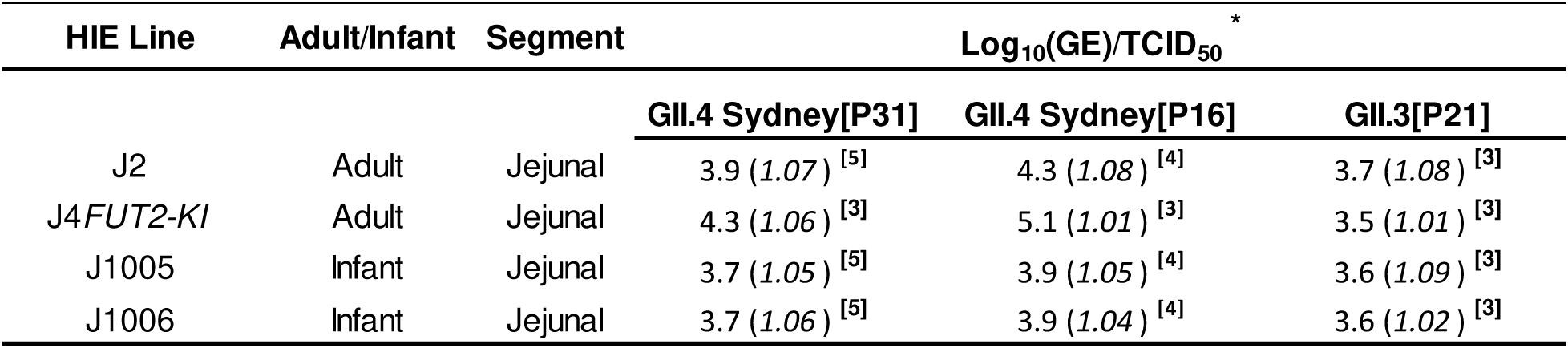
GE/TCID_50_ values of GII.3 and GII.4 in different jejunal HIEs. Differentiated monolayers were inoculated, six wells per dilution in each experiment, with two-fold serial dilutions of GII.4 Sydney[P31], GII.4 Sydney[P16], or GII.3[P21]. After 1 hpi, monolayers were washed twice and cultured in the OGMd medium (+ 500 μM GCDCA). * Values represent geometric mean GE/TCID_50_ from n=3-5 experiments and expressed as log_10_(GE) required to achieve infectivity of 50% of the inoculated cultures at 24 hpi; values in parentheses correspond to SD (standard deviation); superscript **[n]**: number of independent experiments.

Building upon our previous investigation where we demonstrated the successful replication of one GI genotype and 11 GII genotypes using the established HuNoV cultivation system (23), the current study demonstrates the capability of J4*FUT2-KI* to support enhanced replication of GII.4, GII.3, and GII.6 HuNoVs, as indicated by the data presented above. Based on these findings, we aimed to further elucidate the replication of additional HuNoV strains in J4*FUT2-KI* HIE monolayers. At 24 hpi, we observed an increase in viral RNA levels, with a net log_10_ difference in viral growth, ranging between 0.5 and 3.0, compared to viral RNA levels at 1-2 hpi (Table 3). By using this HIE line in our screening, we expanded the spectrum of cultivatable HuNoV strains to encompass three additional GI genotypes, GI.4, GI.5, and GI.7, and an additional GII.4 Hunter 2004 variant (Table 3, new replicating virus strains bolded).

**Table 3:** Cultivated HuNoV genotypes in the J4*FUT2-KI* HIE line. ND, not determined; infections with (^¶^) were performed in in-house media; infections without (^¶^) were performed in OGM media; Bolded strains have not been reported before; all infections were conducted in the presence of GCDCA.

| Genogroup | Genotype | Reference strain | P Type | Cultivable strains | Log<sub>10</sub> increase in viral RNA |
| --- | --- | --- | --- | --- | --- |
| GI | GI.1 | GI.1[P1]/1968/Norwalk | P1 | 1 | 1.9 |
|  | **GI.4** | **PCO-1993** | P4 | 1 | 1.1 |
|  | **GI.5** | **PCO-1638** | P4 | 1 | 0.6 |
|  | **GI.7** | **PCO-475** | ND | 1 | 0.8 |
| GII | GII.1 | TCH18-98 | P16 | 1 | 1.7 |
|  | GII.2 | TCH05-951 | P2 | 1 | 2.1 |
|  | GII.3 | TCH04-577 | P21 | 2 | 3.0 |
|  |  | PCO-2061 | P12 |  | 1.8 |
|  | GII.4 Lanzou_2002 | TCH02-276 | P4 | 1 | 0.8 |
|  | GII.4 Farmington | TCH04-191 | P4 | 2 | 2.3 |
|  |  | TCH02-539 | ND |  | 0.9 |
|  | **GII.4 Hunter_2004** | **TCH05-797** | P4 | 1 | 1.8 |
|  | GII.4 Yerseke_2006a | TCH07-194 | P4 | 3 | 2.0 |
|  |  | TCH02-186 | P4 |  | 1.5<sup>¶</sup> |
|  |  | TCH02-276 | P4 |  | 0.8<sup>¶</sup> |
|  | GII.4 Den Haag_2006b | TCH07-882 | P4 | 6 | 2.2 |
|  |  | MDA09-01 | P4 |  | 2.4 |
|  |  | TCH23-323 | P16 |  | 2.3 |
|  |  | TCH08-227 | P4 |  | 1.5<sup>¶</sup> |
|  |  | TCH08-429 | P4 |  | 1.3<sup>¶</sup> |
|  |  | TCH08-430 | P4 |  | 0.9<sup>¶</sup> |
|  | GII.4 New Orleans_2009 | TCH11-64 | P4 | 2 | 1.7 |
|  |  | TCH23-25 | P4 |  | 0.9 |
|  | GII.4 Sydney_2012 | TCH12-580 | P31 | 7 | 2.7 |
|  |  | TCH14-10 | P31 |  | 2.0<sup>¶</sup> |
|  |  | TCH15-82 | P31 |  | 2.7 |
|  |  | TCH15-88 | P31 |  | 3.0<sup>¶</sup> |
|  |  | TCH15-123 | P16 |  | 2.0<sup>¶</sup> |
|  |  | BCM16-1 | P31 |  | 2.0 |
|  |  | BCM16-16 | P16 |  | 2.2 |
|  | GII.4 Sydney_2015 | BCM16-22 | P16 | 4 | 1.8<sup>¶</sup> |
|  |  | TCH19-76 | P16 |  | 2.3 |
|  |  | BTH18-10 | P16 |  | 2.0 |
|  |  | BTH19-9 | P16 |  | 0.8 |
|  | GII.6 | TCH13-106 | P7 | 4 | 1.3 |
|  |  | TCH15-167 | P7 |  | 2.9 |
|  |  | TCH08-166 | P7 |  | 1.0<sup>¶</sup> |
|  |  | CVID | ND |  | 1.3 |
|  | GII.7 | TCH06-163 | P8 | 1 | 0.5<sup>¶</sup> |
|  | GII.8 | TCH09-279 | P7 | 1 | 0.7 |
|  | GII.12 | TCH09-477 | P8 | 1 | 1.1 |
|  | GII.13 | TCH10-338 | ND | 1 | 0.9 |
|  | GII.14 | TCH14-364 | P16 | 1 | 1.5 |
|  | GII.17 | TCH14-385 | P38 | 2 | 2.1 |
|  |  | 1295-44 | P13 |  | 1.5 |

## Discussion

In our previous work, we reported conditions that enhanced the HIE culture system for HuNoV replication, marked by an expansion in the number of cultivatable strains and increased replication magnitude (23). Since then, we have established several novel HIE lines, prompting us to undertake a new evaluation to ascertain whether any of these newly established lines exhibit better performance for supporting HuNoV replication compared to our existing culture system. Additionally, the introduction of the ODM medium in 2020, by STEMCELL Technologies, raised the question about its potential to outperform the OGM media in supporting HuNoV replication.

Our comparative analysis between OGM and ODM media revealed both media support viral replication, but there was a significant enhancement in the replication efficiency of GII.4 Sydney[P31] and GII.3[P21] with OGM medium. This comparison, not previously undertaken in other studies that used OGM or ODM media (28, 39–43), provides guidance for laboratories using commercial media for HuNoV studies. Further, an evaluation of newly established lines, derived from intestinal segments of different donors, highlights strain-, donor- and segment-specific differences within the GII genogroup of HuNoVs. Varying viral replication efficiency between different intestinal segments and donors, suggests differences in receptor expression, innate immune responses, or other host factors. An unexpected result was complete lack of permissiveness for GII.3 virus replication of HIE cultures from any intestinal segment of the secretor positive 2003 donor while a GII.4 virus replicated normally in the HIEs from the small intestinal segments of this donor. The 2003 line is the first secretor positive HIE line that we have tested that does not support GII.3 replication. Due to the limited number of small intestinal HIEs from 4 donors, we did not specifically evaluate differences in virus yields in HIEs from the distinct small intestinal segments although the data suggest there may be differences. These results support the continued testing of new lines from more donors. Such studies and more detailed studies of the 2003 line may reveal new host factors including factors that affect cell permissiveness that underlie strain-specific differences in susceptibility to HuNoV infection yet are independent of FUT2 expression and may provide new, valuable insights into the biology and pathogenesis of HuNoV infections. The lack of infection in the colonic HIEs from the same donors, where small intestinal cultures support infection, is consistent with previous histological studies that showed no detection of viral capsid protein in colon biopsies from immunocompromised patients with chronic HuNoV infection and from our previous replication studies in colonic HIEs generated from biopsies of different donors (23, 44). These new studies exclude genetics as being the factor for the lack of replication in colonic cultures.

Our studies also demonstrate that H9hESC-derived H9tHIEs support HuNoV replication. The H9hESCs are capable of differentiation into various cell types, including enteric neuronal cells or immune cells (45, 46), offering potential applications for future co-culture studies for a more comprehensive human intestinal model. The functional properties of these H9tHIE cultures may align more closely with those of human intestinal stem cell-derived HIEs, since they support efficient replication of HuNoV strains (Figure 2). Direct comparison of HuNoV replication to J2 HIEs showed that H9tHIEs provided higher viral yields, specifically for GII.3[P21]. Future studies are needed to examine their quantitative susceptibility through a comparative assessment of TCID_50_ values across multiple viral strains.

Additionally, the replication data indicate that the J4*FUT2-KI* line supports better HuNoV replication compared to the other HIE lines tested. The GII.6 CVID patient-derived virus exhibited optimal replication in J4*FUT2-KI*, with minimal observed replication in J2, and no replication observed in J2*STAT1-KO* and the patient’s own HIE line. The lower titer of this virus, compared to GII.6/TCH13-106, might be a factor contributing to its poor replication. Van Kampen et al. also reported no replication of a CVID patient-derived GII.4 in the J2 line (16). It is tempting to speculate that an increased level of HBGA expression accounts for the enhanced ability of the J4*FUT2-KI* HIEs to support better replication of the GII.6 CVID (BCM18-1) or other HuNoVs (Table 3). However, the level of glycan expression in J4*FUT2-KI* HIEs is similar to that in the J2 HIE line (30), so we hypothesize that the improved replication may be linked to other altered host factors such as innate immune responses that remain to be fully understood. Studies with a CVID virus and the CVID HIE line represent opportunities to dissect biologic properties and viral evolution in a chronically infected host.

Our results confirm that infant HIE lines (38) support good GII.3 and GII.4 viral yields but these do not surpass yields in adult J2 and J*4Fut2-KI* HIE lines, as seen with another virus strain in these lines (38). Comparative assessment, based on inoculation of infant and adult HIEs revealed the J4*FUT2-KI* HIE line has a similar susceptibility to GII.3 viral infections as measured by GE/TCID_50_ but infection results in higher virus yields.

Many of our studies that screen new virus strains for positive infectivity simply report the ability of an HIE line to support replication of a specific HuNoV strain in stool. We use, where possible, inocula with input GEs/well above the previously reported minimal dose needed to detect replication in HIEs for most virus strains (47). We are now further determining the GE/TCID_50_ values of each inoculum to obtain comparative quantitative data on the infectivity of each virus strain in each HIE line. Determining the GE/TCID_50_ values for the various virus inocula for the different HIE lines allows comparisons of the permissiveness of the different HIE lines and standardization of infections by using equivalent infectious doses (15, 48). This standardization is crucial because each stool inoculum contains different GEs and likely different particle to infectious particle ratios. This standardization also is crucial in quantitative therapeutic and comparative studies.

In these new studies, the GE/TCID_50_ comparisons led to the unexpected discovery that J4*FUT2-KI* HIEs are more resistant, compared to the J2 HIE line, to infection with two distinct GII.4 Sydney strains (Table 2). While it might be expected that overexpression of FUT2 would increase susceptibility to HuNoV infection, resulting in lower TCID50 values, our findings indicate otherwise. This unexpected result for GII.4 viruses suggests that factors beyond FUT2 expression or levels of HBGA on the cell surface must influence the susceptibility of this cell culture to viral infection. Such host factors may be involved in viral entry, RNA replication or virus assembly in the J4*FUT2-KI* line. Our previous research has highlighted that compared to other HuNoVs whose entry is mediated by bile-acid induced endocytosis, GII.4 strains enter J2 HIEs by unique mechanisms (43, 49, 50). GII.4 entry and replication does not require bile acid but involves membrane wounding, ASM-mediated lysosomal exocytosis, endosomal acidification and clathrin-independent carriers (CLIC) pathways (49), and GII.4 can overcome host interferon-mediated innate responses (51). These pathways may be different in the genetically modified J4*FUT2-KI* cultures, and further investigation is required to fully understand the molecular mechanisms responsible for the higher TCID50 values of GII.4 viruses in J4*FUT2-KI* HIEs.

Overall, the development of reliable cultivation systems for HuNoV has been a significant challenge for over five decades. Despite numerous attempts, many reported cultivation methods faced reproducibility issues (52–57). While some progress has been made in cultivating HuNoV in B cells and zebrafish models, these systems do not provide the breadth of strain coverage needed for comprehensive HuNoV research and they do not fully recapitulate host susceptibility and pathogenesis as recently reviewed (57–61). Establishment of the HIE system has significantly advanced HuNoV research by enabling cultivation of multiple HuNoV genotypes, discovery of strain-specific differences in HuNoV replication, host responses to infection, and allowing evaluation of neutralizing antibodies and therapeutics. In this study, we report successful cultivation of one additional GII and three GI genotypes, representing a substantial increase in the diversity of strains and allowing for more comparative studies and antiviral screening.

In conclusion, this study significantly provides new insight into HuNoV infection by confirming viral replication in diverse HIEs, including H9tHIEs, those from a CVID patient, organ donors, and infants. Incorporating HIEs from multiple donors reveals segment-and strain-specific susceptibility to HuNoV infection. H9tHIEs effectively mimic the human intestinal environment. We have expanded the spectrum of cultivatable

HuNoV strains, demonstrating positive replication for 4 GI and 11 GII genotypes. The J4*FUT-KI* line remains particularly promising for the growth of numerous HuNoV genotypes. Using OGM media and standardizing infections with TCID50 determinants, help ensure consistent and comparable results by providing a more accurate quantitation of the amount of GEs required for infection. These findings collectively contribute to a more robust and standardized approach to HuNoV cultivation, and continue enhancing our understanding of HuNoV biology with implications for antiviral testing and therapeutic discovery.

## Materials and methods

### HIES and viruses

HIE cultures used in this study are from an HIE bank maintained by Gastrointestinal Experimental Model Systems Core (GEMS) of the Texas Medical Center Digestive Diseases Center (TMC DDC) (Table 4). The use of human tissues to establish HIE cultures was approved by the Baylor College of Medicine Institutional Review Board. HIEs were generated from distinct intestinal segments from organ donors provided by the organ donation group LifeGift (Houston, TX, USA). Whole intestines were delivered on ice one hour after arrival (62). The intestinal regions were identified as follows: the duodenum was taken from the first 10 cm of the small intestine; jejunum and ileum were separated from the remaining small intestine using the vascularization of the tissue as a guide; the colon region was identified by the morphology of the colon and the patterning of mesenteric fat. Following identification and separation, the intestinal regions were washed with calcium/magnesium-free PBS buffer. HIE cultures were established from small pieces of each intestinal region as previously described (63, 64). The D_201_HIE line was generated from a duodenal biopsy of an individual with congenital variable immunodeficiency who was chronically infected with human norovirus. H9tHIE line was generated from H9 human Embryonic Stem Cell (H9hESC)-derived HIOs transplanted under kidney capsule of immunocompromised mice. J2*STAT-KO* and J4*FUT2-KI* HIEs are genetically modified and described previously (30). HIEs J1005 and J1006 were generated from two infants, 10 and 12 weeks at surgery (corrected gestational age, 37 and 42 weeks, respectively (65). All HIEs used in this study are secretor positive (see Table 4). Propagation of HIEs as multilobular cultures and preparation of monolayer cultures for infection were performed as previously described (23). Virus inocula were stool filtrates prepared and stored in aliquots at −80°C as previously described (9).

**Table 4:** HIE lines used in this study. Se, secretor; Le, Lewis; y, years; wks, weeks; D, duodenum; J, jejunum, I, ileum; C, colon; (^●^): H9tHIE were generated from 8-week-old tHIOs which are developmentally similar to an 18 gestational week human fetal intestine (36); (^●●^): (age at surgery/corrected gestational age at surgery).

| HIE line | Source age | Phenotyping results |  |  |
| --- | --- | --- | --- | --- |
|  |  | Secretor<br>Status | ABH<br>Status | Lewis<br>Status |
| J2 | 52 y | Se+ | B | Le<sup>b</sup> |
| J4<sup>FUT2/KI</sup> | 62 y | Se+ | O | Le<sup>b</sup> |
| LG2002 (D,J,I,C) | 34 y | Se+ | O | Le<sup>b</sup> |
| LG2003 (D,J,I,C) | 23 y | Se+ | O | Le<sup>b</sup> |
| LG2004 (D,J,I,C) | 25 y | Se+ | A | Le<sup>b</sup> |
| LG2005 (D,J,I,C) | 23 y | Se+ | O | Le<sup>b</sup> |
| H9tHIE | 18 wks<sup>•</sup> | Se+ | B | Le<sup>b</sup> |
| J1005 | 10/37 wks<sup>••</sup> | Se+ | A | Le<sup>b</sup> |
| J1006 | 12/43 wks<sup>••</sup> | Se+ | B | Le<sup>b</sup> |
| D201 | 56 y | Se+ | O | Le<sup>b</sup> |

### Media

Different media were used to maintain and differentiate HIEs:

1. A complete medium with growth factors (WRN medium), prepared at BCM by the DDC core, consisted of CMGF[−] medium (46%; v/v) (9) supplemented with mouse recombinant epidermal growth factor (EGF, 50 ng/mL final concentration; Invitrogen), nicotinamide (10 mM; Sigma), gastrin I (10 nM; Sigma), A-83-01 (500 nM; Tocris), SB202190 (10 uM; Sigma), B27 supplement (1X; Invitrogen), N2 supplement (1X; Invitrogen), *N*-acetylcysteine (1 mM; Sigma), and the conditioned medium (50%; v/v) prepared from L-WRN cell line (ATCC CRL-3276) that co-expresses Noggin, R-spondin, and Wnt-3A growth factors.
2. Commercial Intesticult human organoid growth medium (Stem Cell Technologies; Cat#06010) is composed of two components, the OGM human basal medium and the organoid supplement. The cell pellets, resulting from HIE cell dispersion, were suspended in the proliferation OGM medium (OGMp), prepared by mixing equal volumes of basal medium and organoid supplement, and supplemented with 10 μM ROCK inhibitor Y-27632.
3. After 1 day of cell growth as a monolayer, the OGMp medium was changed to the differentiation OGM (OGMd), consisting of an equal volume of basal medium and CMGF[−] medium. The cell monolayers were differentiated for 5 days as previously described.
4. The commercial Intesticult human differentiation medium (ODM, Stem Cell Technologies; Cat# 100-0214) is another medium used to differentiate HIE monolayers.

### Viral infection

Five-day differentiated HIE monolayers were washed once with CMGF(−) medium and inoculated with HuNoV for 1-2 hr at 37°C. The inoculum was removed, and monolayers were washed twice with CMGF(−) medium to remove the unbound virus. OGMd or ODM differentiation medium [100 μL containing 500 μM bile acid glycochenodeoxycholic acid (GCDCA)] was then added, and the cultures were incubated at 37°C for the indicated time points.

### Tissue culture infectious dose 50% (TCID_50_) assay

We determined GE/TCID_50_ values to allow us to evaluate the permissiveness of each HIE line for infection with each virus and we previously reported that HuNoV strains have different GE/TCID_50_ values for each HIE line (15, 48). We determined the TCID_50_ per inoculum by examining the numbers of wells that show increases in genomic equivalents of virus per well at 24 hr above baseline (at 1 hr) as measured by RT-qPCR (15, 48). TCID50 per volume were calculated by Reed-Muench. We also used RT-qPCR to measure the number of GEs per volume using a genogroup-specific standard. We then calculated the GE per TCID50. Compiled data from 3-5 independent experiments are presented. In this context, the information is expressed as the number of GEs [log_10_(GE)] per TCID50) at which 50% of the cultures are infected as determined by RT-qPCR.

Four HIE lines from 2 infants (J1005 and J1006) and 2 adults (J4*FUT2-KI*, and the prototype J2) were seeded as monolayers in 96-well plate for 24 hrs in OGM proliferation medium supplemented with 10 μM Rock inhibitor Y-27632, and differentiated for 5 days in OGMd medium by changing the medium every other day. Prior to infection, one aliquot of the GII.4 Sydney[P31] (1.8×10^7^ GEs/μL), GII.4 Sydney[P16] (4.3×10^6^ GEs/μL), or GII.3 (7.4×10^6^ GEs/μL) stool samples was thawed and diluted 1:5,000 in Dulbecco’s phosphate buffer saline calcium/magnesium free (DPBS; ThermoFisher). This viral dilution was 2-fold serially diluted in CMGF(-) basal medium supplemented with 500 μM GCDCA bile acid in a 96-well round-bottom plate. Six replicates were typically prepared for each dilution. HIE monolayers were then infected with 100 μL of each dilution for 1 hr at 37°C under 5% CO_2_ atmosphere, washed twice with CMGF(-) basal medium, and incubated in 100 μL/well OGMd with 500 μM GCDCA. After 24 hpi, total RNA was extracted from each well and viral replication was assessed by RT-qPCR in duplicate.

### Quantification of viral replication by RT-qPCR

RNA extraction and RT-qPCR were performed as previously described (23). In brief, total RNA was extracted from each infected well using the KingFisher Flex purification system and MagMAX-96 Viral RNA isolation kit. RNA extracted at 1 hpi was used as a baseline to determine the amount of input virus that remained associated with cells after washing the infected cultures to remove unbound virus. The primer pair and probe COG2R/QNIF2d/QNIFS (66) were used in the RT-qPCR for detection of GII genotypes and the primer pair and probe NIFG1F/V1LCR/NIFG1P (67) were used for GI.1. RT-qPCR was performed with qScript XLT One-Step RT-qPCR ToughMix reagent with ROX reference dye (Quanta Biosciences) in an Applied Biosystems StepOnePlus thermocycler. Each extracted RNA was run in duplicate with the cycling conditions: 50°C (15 min), 95°C (5 min), followed by 40 cycles of 95°C (15 s) and 60°C (35 s). Standard curves based on recombinant GII HuNoV RNA transcripts were used to quantitate viral genome equivalents (GEs) in RNA samples. The limit of detection of the RT-qPCR assay was 20 GEs. A threshold for successful viral replication was established by considering a 0.5 increase in log_10_(GE) after 24 hpi relative to the genomic RNA detected at 1 hpi (23).

### Statistical analysis

Each experiment was performed more than once, with three technical replicates of each culture condition and time point. Compiled data from at least two experiments was presented. All statistical analyses were performed on GraphPad Prism version 10.2.3 for Windows (GraphPad Software, La Jolla, CA, USA). Comparison between groups was performed using the one-way ANOVA with Tukey’s multiple comparisons test or Student’s t-test, which was applied to the data presented in Figure 1. *P* values of <0.05 were considered statistically significant.

## Data availability

All data are available in the paper. All HIE cultures used in this study are from an HIE bank maintained by the Texas Medical Center Digestive Diseases Center (TMC DDC) core https://www.bcm.edu/research/research-centers/texas-medical-center-digestive-diseases-center.

## Acknowledgements

This work was funded in part by Public Health Service grant P01-AI057788 to M.K.E, R.L.A., and BVV Prasad, P30 DK 056338 (to H. El-Serag), which supports the Texas Medical Center Digestive Diseases Center and GEMS Core, and NIH 1K08DK131326 to A.L.S. The authors acknowledge the Advanced Technology Core Laboratories (Baylor College of Medicine), specifically the Integrated Microscopy Core with funding from the NIH (DK56338, CA125123, ES030285), and 398 CPRIT (RP150578, RP170719). We thank LifeGift, Javier Nieto, and the donor families for providing intestinal tissues for generating HIEs.

## Notes

### Competing Interest Statement

The authors have declared no competing interest.

### Summary of Updates

This version of the manuscript was updated to respond to the reviewers comments. Changes were made to the figures, the text, as well as the title.

